# Chromosome-level assembly of the common lizard *(Zootoca vivipara)* genome

**DOI:** 10.1101/520528

**Authors:** Andrey A. Yurchenko, Hans Recknagel, Kathryn R. Elmer

## Abstract

Squamate reptiles exhibit high variation in their traits and geographical distribution and are therefore fascinating taxa for evolutionary and ecological research. However, high-quality genomic recourses are very limited for this group of species, which inhibits some research efforts. To address this gap, we assembled a high-quality genome of the common lizard *Zootoca vivipara* (Lacertidae) using a combination of high coverage Illumina (shotgun and mate-pair) and PacBio sequence data, with RNAseq data and genetic linkage maps. The 1.46 Gbp genome assembly has scaffold N50 of 11.52 Mbp with N50 contig size of 220.4 Kbp and only 2.96% gaps. A BUSCO analysis indicates that 97.7% of the single-copy Tetrapoda orthologs were recovered in the assembly. In total 19,829 gene models were annotated in the genome using a combination of three *ab initio* and homology-based methods. To improve the chromosome-level assembly, we generated a high-density linkage map from wild-caught families and developed a novel analytical pipeline to accommodate multiple paternity and unknown father genotypes. We successfully anchored and oriented almost 90% of the genome on 19 linkage groups. This annotated and oriented chromosome-level reference genome represents a valuable resource to facilitate evolutionary studies in squamate reptiles.

## 1. BACKGROUND

Squamate reptiles (lizards and snakes) are one of the biggest groups of vertebrate animals, with more than ten thousand species distributed worldwide (Uetz and Hošek, 2015). Squamates exhibit fascinating variation in their morphology, development, reproductive strategies, sex determination mechanisms, karyotype and environmental adaptations (Sites et al. 2011). The major lineages of Squamata diverged around 100-200 million years ago (Zheng and Wiens, 2016; Irisarri et al. 2017) forming an incredible diversity of species, much of which remains cryptic and under-studied (Pinho et al. 2007; Oliver et al. 2009; Guarnizo et al. 2016). Squamate reptiles show an extraordinary frequency of evolving complex biological traits, such as egg-laying and live-bearing (Blackburn 2006; Pyron and Burbrink 2014; King and Lee 2015), parthenogenesis (Neaves and Baumann 2011; Sitres et al. 2011), and a high variability of karyotypes including the presence of both micro and macro chromosomes (Deakin and Ezaz 2014).

The recent technological advances in genome sequencing and assembly methods enabled the first squamate genome to be sequenced in 2011; the famous ecological and evolutionary model species the green anole lizard (Alföldi et al. 2011). A modest 60% of the available anole genome is placed to chromosomes. Similarly, other recently published squamate genome assemblies are assembled into scaffolds (e.g. Castoe et al. 2013; Liu et al. 2015) but often are fragmented (Kolora et al. 2018; Tollis et al. 2018). The recent publication of the wall lizard genome marks the first step towards high-quality chromosome-level assemblies in squamate reptiles (Andrade et al. 2018). However, the previous lack of high-quality genomic assemblies for this important group of animals has slowed research on chromosomal evolution, environmental adaptation, the genomic basis of venom evolution, and other important questions of evolutionary biology. With the development of increasingly economical sequencing technologies and standardized methodological approaches, more genomes will become accessible in the near future.

In this study we combined high-coverage Illumina-derived sequencing with multilayer PacBio and RNA-seq based scaffolding to generate a high-quality genome assembly of the Eurasian common lizard, *Zootoca vivipara* (Lacertidae). During the genome assembly process, we extensively validated and improved the assembled scaffolds with a newly generated, ddRADseq-based high-density linkage map produced with an original methodology from wild-caught samples. This combined assembly strategy allowed us to place nearly 90% of the 1.46 Gbp genome to 19 linkage groups and produce an overall high-quality assembly, with 97.7% of the Tetrapoda-specific single-copy orthologs recovered. *Zootoca vivipara* is a fascinating ecological and evolutionary model and is the focus of considerable research; it has the broadest natural range among vertebrates, the most northern distribution for reptiles, several major lineages with a divergence time of maximally ca. 6 million years (Surget-Groba et al. 2001; Recknagel et al. 2018), and striking differences in reproductive mode (viviparous and oviparous) and sex chromosome karyotypes (Odierna et al. 2004; Kupriyanova et al. 2008). An available reference genome of this lizard will facilitate studies of parity evolution, chromosomal architecture of sex determination, and environmental adaptations exhibited by this and other squamate reptiles.

## 2 RESULTS

### 2.1 Genome assembly

The estimated genome size of *Z. vivipara* was ≈ 1.345 Gbp (2.914 Gbp for human in the same estimation) based on SGA k-mer distribution analysis and agreeing with earlier flow-cytometry based reports that estimated the genome size to be in range between 1.035 Gbp and 1.515 Gbp (Desmet 1981, Vinogradov 1998).

The pipeline for our assembly involved combining long- and short-read data from DNA sequencing, and gap-closing with long-read DNA sequence and transcriptome sequence (Fig. 1). This genome sequence was then anchored and oriented with species-specific linkage maps (see below).

**Fig. 1.**
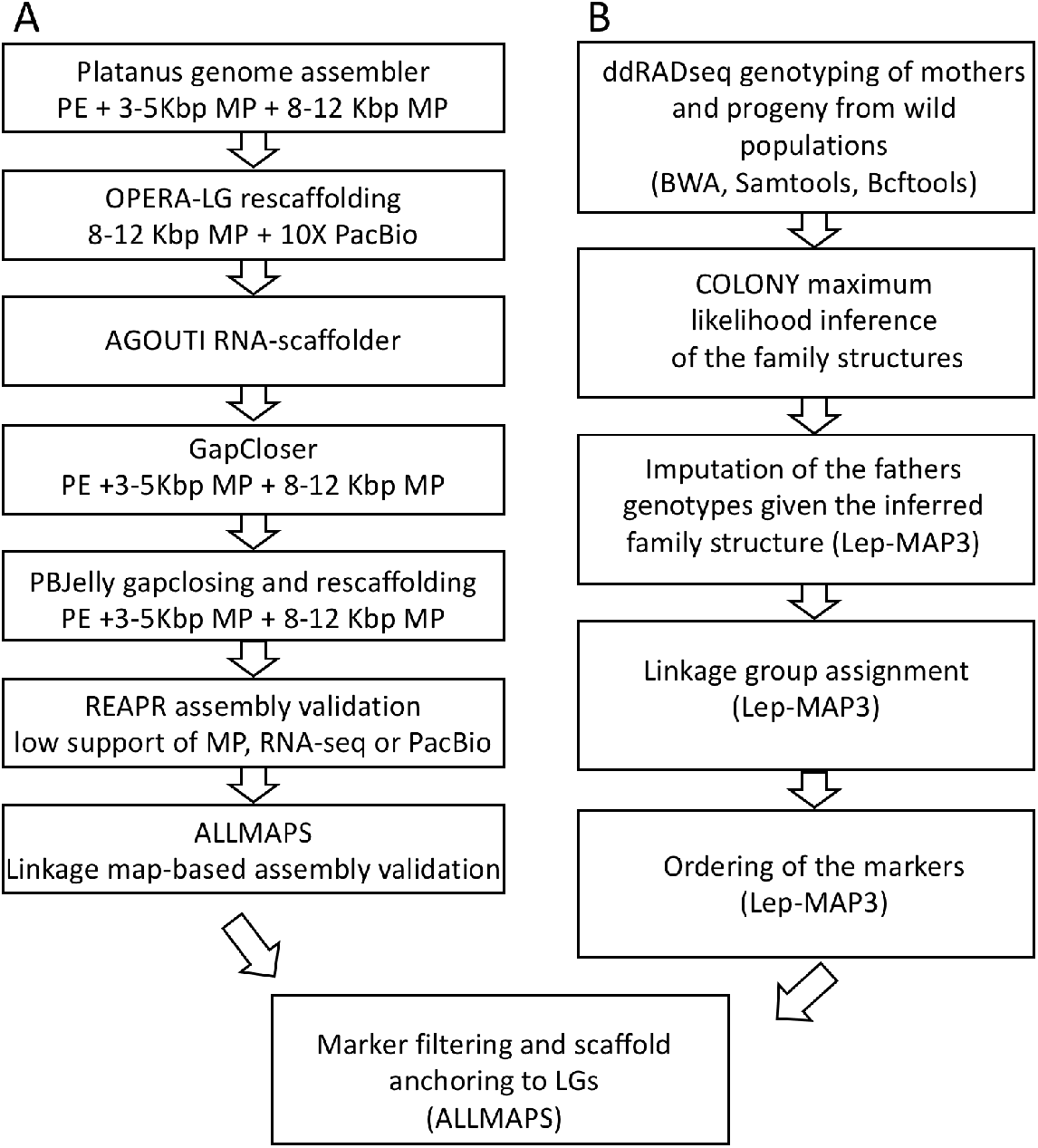
The genome assembly (A) and linkage map generation (B) pipelines used in the study

For the genomic short-read illumina data, after data filtering and read correction we received 343 M shotgun paired-end (PE) reads, 78 M reads from 3-5 Kbp mate-pair libraries, and 53 M reads from 8-12 Kbp mate-pair libraries. These were used to build contigs and scaffolds using the Platanus genome assembler (Table 1). Additionally, we used 102 M paired-end and 164 M single-end reads that were a by-product after trimming and filtration of the PE and mate-pairs. These reads were used only for the initial contig assembly.

**Table 1.** Sequencing used for the assembly. The total data for genome assembly (excluding linkage maps) was 741 M reads of DNA sequence and 195 M reads of RNA sequence.

| **Data type** | **Before filtering, million reads** | **After filtering, million reads** |
| --- | --- | --- |
| Illumina shotgun sequencing, PE 150 bp | 394 | 343 |
| 3-5 kbp mate-pairs, PE 150 bp | 197 | 78 |
| 8-12 kbp mate-pairs, PE 150 bp | 140 | 53 |
| PacBio (N50 11498 kbp) | 2,1 | 1,7 |
| RNA-seq, PE 150 bp | 357 | 195 |
| RAD-seq, PE 150 bp for the linkage map | 763 | 643 |
| SE from mate-pairs and shotgun sequencing | - | 164 |
| PE from mate-pairs and shotgun sequencing | - | 102 |

The first scaffolds produced with Platanus assembler had N50 metrics equal to 5.35 Mbp and consisted of 366.9k contigs with a N50 of 5.23 Kbp (Table 2). At the next step, we re-scaffolded the assembly with Opera-LG using PacBio (1.7 million reads) and 8-12 Kbp mate-pairs, which allowed us to double the N50 scaffold length (12.52 Mbp). The next steps were re-scaffolding with RNA-seq information about splicing events, and gap closing using short reads; these steps increased the N50 contig size to 83.4 Kbp. Further gap closing with long PacBio reads allowed us to additionally increase the contig length distribution size to achieve N50 of 220.4 Kbp.

**Table 2.**
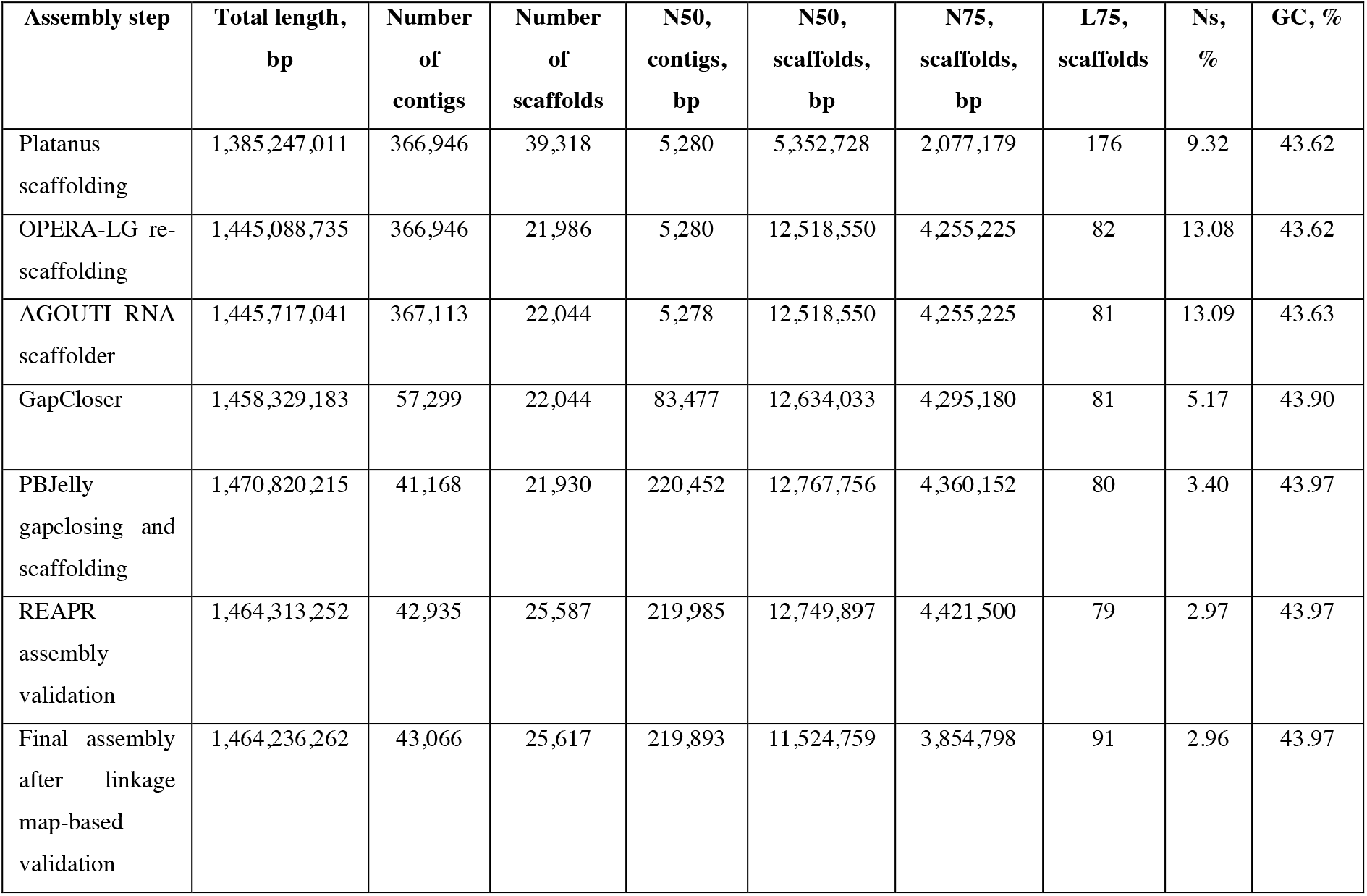
The main steps and associated statistics of the genome assembly

| Assembly step | Total length, bp | Number of contigs | Number of scaffolds | N50, contigs, bp | N50, scaffolds, bp | N75, scaffolds, bp | L75, scaffolds | Ns, % | GC, % |
| --- | --- | --- | --- | --- | --- | --- | --- | --- | --- |
| Platanus scaffolding | 1,385,247,011 | 366,946 | 39,318 | 5,280 | 5,352,728 | 2,077,179 | 176 | 9.32 | 43.62 |
| OPERA-LG re-scaffolding | 1,445,088,735 | 366,946 | 21,986 | 5,280 | 12,518,550 | 4,255,225 | 82 | 13.08 | 43.62 |
| AGOUTI RNA scaffolder | 1,445,717,041 | 367,113 | 22,044 | 5,278 | 12,518,550 | 4,255,225 | 81 | 13.09 | 43.63 |
| GapCloser | 1,458,329,183 | 57,299 | 22,044 | 83,477 | 12,634,033 | 4,295,180 | 81 | 5.17 | 43.90 |
| PBJelly gapclosing and scaffolding | 1,470,820,215 | 41,168 | 21,930 | 220,452 | 12,767,756 | 4,360,152 | 80 | 3.40 | 43.97 |
| REAPR assembly validation | 1,464,313,252 | 42,935 | 25,587 | 219,985 | 12,749,897 | 4,421,500 | 79 | 2.97 | 43.97 |
| Final assembly after linkage map-based validation | 1,464,236,262 | 43,066 | 25,617 | 219,893 | 11,524,759 | 3,854,798 | 91 | 2.96 | 43.97 |

At the next step we validated the genomic assembly to reduce the number of possible misjoinings. The REAPR pipeline customised with additional PacBio and RNA-seq data allowed us to identify 1733 likely erroneous joins between contigs, mostly at the ends of scaffolds that were then broken further.

At the final stage of our workflow we identified 30 intrascaffold regions that showed signs of misassembly according to the linkage map data (see **2.2 Linkage map construction**). Linkage maps cannot locate exact breakpoints of misassembly but approximate them very well (Tang et al. 2015), thus we used this information to break the assembly and to remove suspicious joinings. After this validation step, the formal assembly quality metrics slightly reduced (scaffold N50 by 1.23 Mbp to 11.52 Mbp), but still indicated a high level of assembly contingency.

### 2.2 Linkage map construction

For the linkage map construction, in total 643M clean PE reads were used representing data from 205 individuals in 20 families of mothers and offspring. Estimation of family structure and parent assignment demonstrated widespread multiple paternity in Z. *vivipara*. Four of 20 families had a single father while all other families had from two to four fathers, with the mean number of progeny per half-sib family equal to 3.7. The assignment of progeny to different fathers was additionally supported by an exploratory PCA analysis of each family.

The genotype calling resulted in a total of 1,442,510 unfiltered SNPs for all the individuals. After filtering we retained 109,640 high-quality biallelic SNPs and used them for imputing the missing genotypes of fathers. At the first stage of linkage map construction, 17,210 markers were assigned to 19 linkage groups (from 395 to 1648 markers per LG, LOD score=10.7), which corresponds to the *Z. vivipara* karyotype with 17 autosomes and the Z and W sex chromosomes (2n=36 chromosomes including ZZ/Zw sex chromosomes) specific for this population (Kupriyanova et al. 2014). At the next step, an additional 7,177 markers were assigned to these LGs with minimal LOD score equal to 9. The ordering of markers was the final stage of the linkage map construction and allowed us to create male and female linkage maps with the entire size equal to 1929.24 and 2263.13 cM respectively. There were 1.27 and 1.24 markers per cM for the male and female linkage maps with 2487 and 2845 unique points for each of them (Table 3).

**Table 3.**
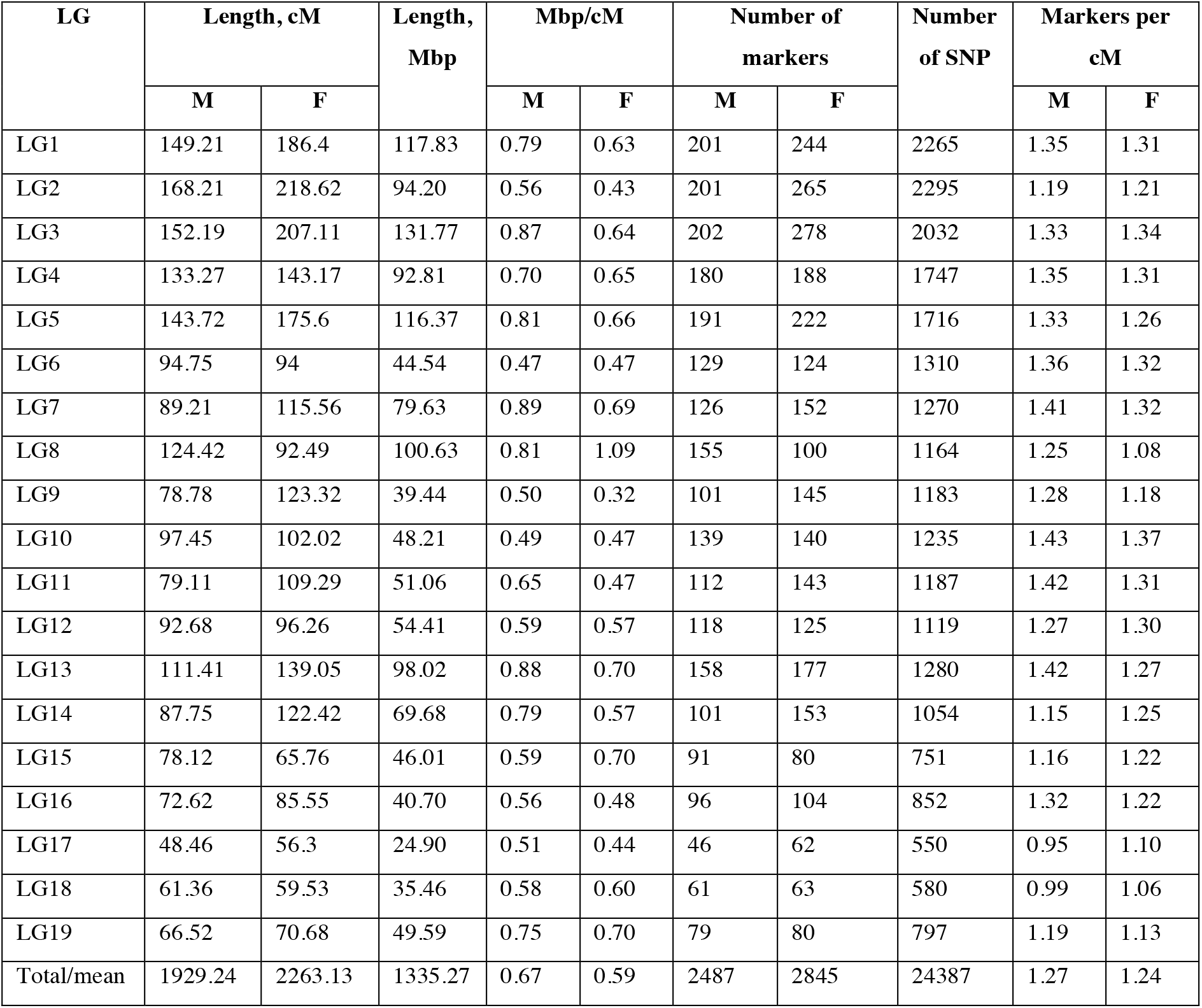
Statistics for the *Zootoca vivipara* linkage map and the anchored assembly

### 2.3 Scaffolding with linkage groups

We anchored scaffolds to the linkage groups using ALLMAPS software and determined that in total 91.2% of the assembly anchored and 89.5% of the assembly oriented using the linkage map (Table 4). We had on average 18.1 SNP markers per Mb of genome, with 286 scaffolds of the 420 scaffolds anchored having more than four markers. The physical size of the linkage groups varied from 24.9 to 131.77 Mbp and physical positions of markers strongly correlated with the linkage-based positions on the map (Figure 2). The average resolution of the male and female linkage maps was 0.67 and 0.59 Mbp per cM respectively (Table 3).

**Figure 2.**
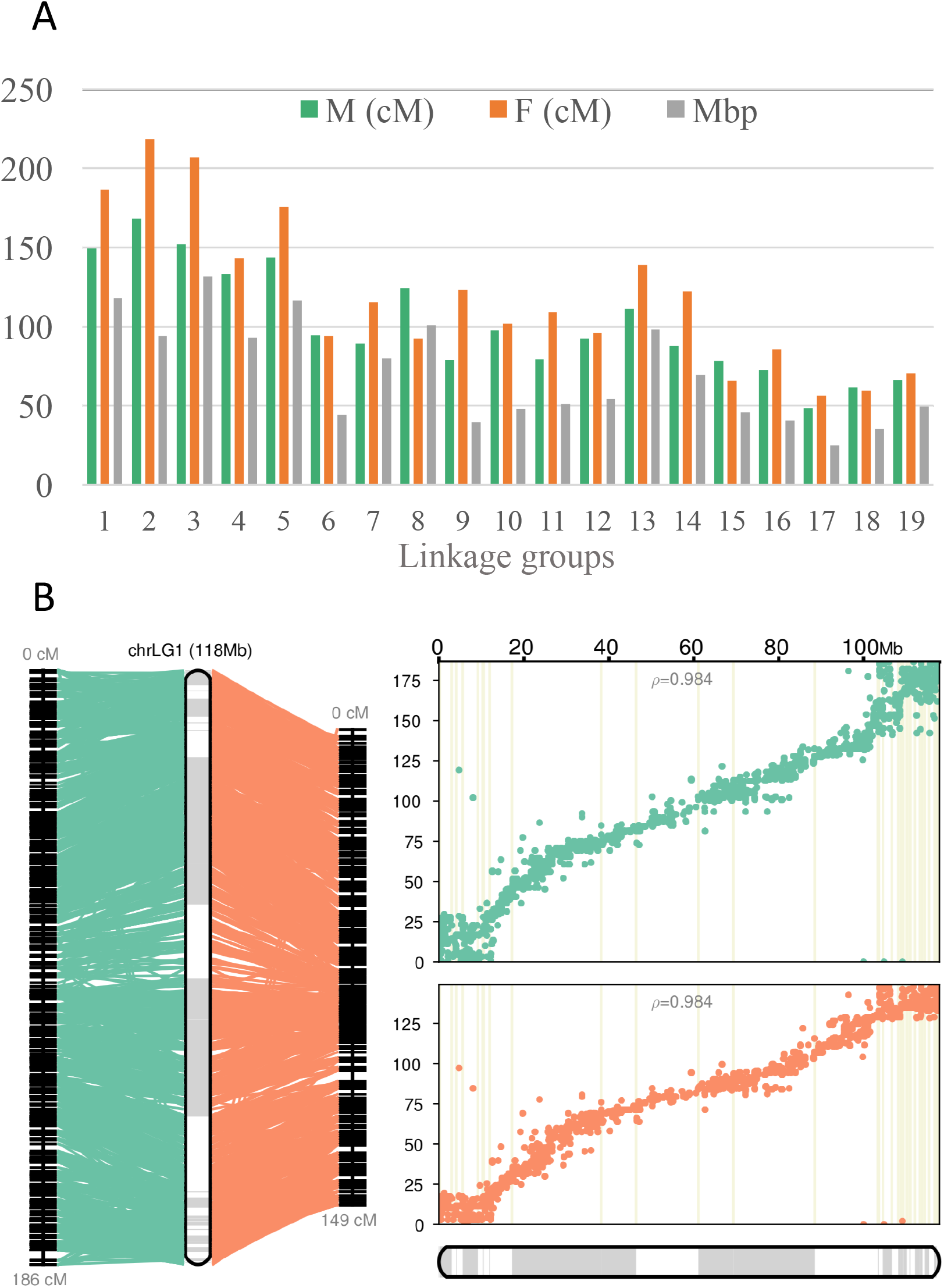
Correlation between genetic and physical size in the common lizard, *Zootoca vivipara*, genome. A) The length of the male (M), female (F) and consensus linkage groups. B) An example of Linkage Group 1 based on the male (green) and female (orange) genetic maps. Pearson correlation coefficients between the physical (X axis) and genetic (Y axis) distances are indicated (*ρ* = 0.984). Grey and white bars represent scaffolds.

**Table 4.** Linkage map summary and association with scaffolds

| Statistics | Anchored | Oriented | Unplaced |
| --- | --- | --- | --- |
| Number of markers | 24,205 | 23,781 | 126 |
| Number of markers per Mbp | 18.1 | 18.1 | 1 |
| Scaffolds | 420 | 295 | 23,235 |
| Scaffolds with 1 marker | 60 | 0 | 38 |
| Scaffolds with 2 markers | 41 | 31 | 14 |
| Scaffolds with $\geq 4$ markers | 286 | 256 | 8 |
| Total bases | 1,335,231,344 (91.2%) | 1,310,909,853 (89.5%) | 129,018,870 (8.8%) |

### 2.4 Genome annotation

To produce a high-quality gene annotation of the *Z. vivipara* genome, we combined *ab initio* prediction with RNA-seq supported homology-based methods. The homology-based method, using GeMoMa, allowed us to identify 21,187 high quality gene-models with strong homology to chicken, Japanese gecko, and anole lizard genomes. We considered these models as the most trustworthy datasets because homology supported by RNA-seq evidence relative to related species is usually highly reliable (Keilwagen et al. 2016). At the same time, to find genes with low homology level to the existing genomes, we relied on the *ab initio* gene prediction, which is very sensitive to gene-like structures in genomes. The *ab initio* AUGUSTUS pipeline identified 15,637 gene-models which were finally combined using EVidenceModeler with 28,473 RNA-seq based Transdecoder and GeMoMa gene models. After filtering out genes without any detected homology to the Swiss-Prot database we received a final set of 19,829 protein-coding gene models.

### 2.5 Genome completeness

In order to further quantify the quality of the final scaffolds we estimated the number of the recovered Tetrapoda single copy orthologues (BUSCO) in the assembled genome and found that 94% of orthologues were completely assembled (with 1.3% of them being duplicated), 3.7% were fragmented, and 2.3% of the 3950 benchmarked genes were missed. This metric indicates that the assembly was of high quality with only minor parts of the genome being fragmented.

## 3. CONCLUSIONS

Here we report the chromosome-level genome assembly of the common lizard, *Z. vivipara*, which is one of the most widely distributed vertebrates in the world and possesses several distinctive biological features such as reproductive bimodality, sex chromosome variation, colour polymorphisms, and broad climatic tolerance (Vercken et al., 2007; Kupriyanova et al., 2014; Horreo et al., 2018; Recknagel et al. 2018). The final genome assembly contains 19 linkage groups with almost 90% of the genome anchored and oriented. The assembly length is 1.46 Gbp, which is close to the previous cytometry-based estimations and our k-mer based estimation. In total, we annotated 19,331 protein coding genes and infer satisfactory BUSCO metrics, with the 97.7% of tetrapoda-specific single-copy orthologues recovered (only 3.7% of them are fragmented). To produce the linkage map we applied a novel pipeline to infer the absent paternity genotypes in family structures complicated by complex reproductive behaviour of reptiles. This reference genome assembly will be a useful resource for a wide range of studies aiming to unravel the genomic basis of fascinating evolutionary diversity of squamate reptiles.

## 4 METHODS

### 4.1 Genome assembly

#### 4.1.1 Short read DNA sequencing and Quality Control

For the Illumina sequencing, high molecular weight DNA was extracted from the tissue of a wild caught individual from Scotland with the Dneasy Blood and Tissue Kit (Qiagen) following the manufacturer’s protocol with additional Riboshedder and phenol-chloroform clean-up. DNA quality was checked using a Qubit quantitation, Nanodrop spectrophotometery, and visualisation on agarose. Concentration was 130 ng/ul and 260/280 was 1.88. A TruSeq PCR-free library with 350 bp insert size was generated by Edinburgh Genomics for shotgun sequencing one run of Illumina HiSeqX. A 3-5 Kbp and a 8-12 Kbp Nextera mate-pair library were generated by Liverpool Centre for Genomic Research and sequenced on one lane of HiSeq4000 at Edinburgh Genomics. This generated a total of 1088 M 150 bp paired end reads.

The raw reads were checked using FastQC v0.11.5 software (Andrews 2010) and then reads with a short-insert size were trimmed, low-quality bases removed, and any remaining parts of sequencing adapters trimmed out using Trimmomatic v036 software (Bolger et al. 2014) with the following settings: remove adapters:2:30:10, LEADING:20, TRAILING:20, SLIDINGWINDOW:4:22, MINLEN:35.

To reduce sequencing errors, we applied a read error correction using QuorUM v.1.1.0 software (Marçais et al. 2015) to the short-insert size (350 bp) paired-ends with the following settings: −s 4G −q 33.

The Nextera junction adapters in the long insert size mate-pairs (3-5, 8-12 Kbp) were removed with NxTrim v0.4.1 program (O’Connel et al. 2015) with the --separate and −l 25 settings. Then we used Trimmomatic v036 to remove adapters and technical sequences in the mate-pair reads with the following settings: remove adapters:2:30:10, LEADING:18, TRAILING:18, SLIDINGWINDOW:4:17, MINLEN:25.

Following quality control and trimming, we estimated expected genome size, relative heterozygosity level, repeat content and sequencing error-rate to assess overall characteristics of the data exploiting cleaned short insert size reads with SGA v0.10.15 tool (Simpson 2014).

#### 4.1.2 PacBio long-read sequencing

To receive high weight molecular DNA, we used a standard phenol-chloroform isolation method (Sambrook and Russel 2006) with minimal shaking. A 20 Kbp insert library was generated by Centre for Genomic Research (University of Liverpool) and sequenced with four cells on a PacBio Sequel at the facility.

#### 4.1.3 RNA-sequencing

Total RNA was extracted from RNAlater-preserved tissue (intestine, lungs, liver, muscle) of the same individual used for the genomic sequencing using PureLink RNA Mini Kits (Life Technologies, Carlsbad, CA), following an adapted protocol from Gunter et al. (2013). RNA samples were quantified with a Qubit 2.0 fluorometer (Life Technologies, Carlsbad, CA) and quality was assessed with a 2200 Tapestation (Agilent, Santa Clara, CA). The libraries were prepared with Illumina TruSeq Total Stranded RNA-seq protocol and sequenced together on one lane of Illumina HiSeq4000 (150 bp paired-ends).

#### 4.1.4 Genome assembly

Contigs were assembled using Platanus v1.2.4 assembler (Kajitani et al. 2014), with default parameters, using short-insert size reads and additional short-insert size reads received by-product from mate-pair libraries. Initial scaffolding was performed using the *platanus scaffold* command with all the reads excluding by-product PE and SE from mate-pair libraries because of variable insert sizes. Next the resulting scaffolds were re-scaffolded with the PacBio long reads (at least 1000 bp long to reduce the chimera rate) and the 8-12 kbp mate pairs using OPERA-LG v2.0.6 software (Gao et al. 2016) using a k-mer size=50.

The scaffolds outputted by OPERA-LG were additionally scaffolded using AGOUTI v0.3.3 software (Zhang et al. 2016), which uses RNA-seq data to join scaffolds and contigs based on spliced exonic reads. To apply the AGOUTI software algorithm, we first identified coding sequences in the draft genome using the AUGUSTUS v3.3 package (Stanke et al. 2004) and mapped the RNA-seq reads to the genome with the BWA v0.7.15-r1140 *mem* algorithm (Li and Durbin 2009). Then AGOUTI was used to calculate the most reliable potential joinings between coding exons using reads with mapping quality of at least 50 (-minMQ 50), no more than 2% of mismatches between the reads and the genome allowed (-maxFracMM 0.02), and having at least 40 links between the joined contigs (-k 40).

At the final stage of the assembly process we closed gaps in the assembly, first using the GapCloser v1.12 module of the SoapDenovo 2 package (Luo et al. 2012) with all the available Illumina reads and then with the PBJelly v15.8.24 (English et al. 2012) long-read based algorithm (- minMatch 8 -minPctIdentity 70 -bestn 1 -nCandidates 20 -maxScore −500 -nproc 53 - noSplitSubreads). The PacBio reads were error-corrected using Canu v1.5 software (Koren et al. 2017) prior to gap-closing with the following settings: corErrorRate=0.15, genomeSize=1400m.

### 4.2 Linkage map

#### 4.2.1 Sequencing and SNP calling the ddRADSeq data

In total 205 individuals from 20 families, which contained mothers and progeny without paternal data, were sampled from the Gailtal region in Austria. DNA was extracted from tissue using the Dneasy Blood and Tissue Kit (Qiagen) following the manufacturer’s protocol. Two genomic libraries were constructed using double-digest restriction-site associated DNA sequencing (ddRADSeq, Peterson et al. 2012) following methods outlined in Recknagel et al. (2015) and modified for illumina. Briefly, 1 ug of DNA was digested using restriction enzymes PstI-HF and MspI and subsequently cleaned with the Enzyme Reaction Cleanup kit (Qiagen). The amount of DNA in each offspring individual was then normalized to the sample with the lowest concentration within a library (275 ng in both libraries) to minimize coverage variation. DNA input for mothers was three times that for offspring (750 ng) to ensure higher coverage and therefore high confidence for the maternal genotype. Illumina specific P1 and P2 adapters were ligated to the sticky ends generated by the restriction enzymes. The ligated DNA fragments were then multiplexed and size-selected using a Pippin Prep (Sage Science) for a ‘tight’ range of 150 – 210 bp. Seven separate PCR reactions (for details see Recknagel et al., 2015) were performed per library and combined (Peterson et al. 2012). Following PCR purification, libraries were electrophoresed on a 1.25% agarose gel, visualised with SYBRSafe (Life Technologies), and bands cut out manually to remove any remaining adapter dimers and fragments outside the size range. Product was extracted from the matrix using a MinElute Gel Extraction Kit (Qiagen). DNA libraries were then quantified using a Qubit Fluorometer with the dsDNA BR Assay and quality and quantity assessed using a TapeStation or Bioanalyzer (Agilent Technologies). The libraries were sequenced at Edinburgh Genomics on two lanes of Illumina HiSeq4000 with 150 bp paired-end reads.

We checked the quality of the raw reads using FASTQC software and removed low-quality reads and the right-end 50 bp of overlapping and technical sequences with Trimmomatic v0.36 (CROP:100 SLIDINGWINDOW:4:19). The cleaned reads were demultiplexed using Stacks v1.46 (Catchen et al. 2011) pipeline with command *process_radtags* (-c -q -r --inline_inline --renz_1 pstI -- renz_2 mspI -i gzfastq --len_limit 5).

We aligned the ddRADSeq reads to the *Z. vivipara* scaffolds with BWA mem software and prepared the sorted BAM files with Samtools. Following the alignment, we called SNPs in all samples simultaneously using *samtools mpileup | bcftools call* pipeline. For the *samtools mpileup* command, we only used reads with mapping quality equal to at least 45 (-q 45) and a minimal base quality equal to 20 (-Q 20) along with other options suitable for the ddRADSeq data (--count-orphans --ignore-overlaps -I -P illumine). For the *bcftools call* command we used -m flag, which invokes multi-sample calling algorithm.

#### 4.2.2 Family assignment

It is known that females of many reptile species, including *Zootoca vivipara*, can produce progeny from different males in within a single clutch (Laloi et al. 2004; Uller and Olsson 2008) and therefore we expected some families in our sample to have multiple fathers. In order to address this issue during linkage map construction and imputation of father genotypes, we used COLONY v2.0.6.3 (Jones and Wang 2010) to assign offspring from each clutch (=family) to half-sib families if they were inferred to have different fathers. We split the raw VCF file for linkage map construction by families using vcftools v0.1.15 and applied the following filters: minimal quality of a variant=700 (--minQ 700), minimal quality of a genotype=20 (--minGQ 20), only biallelic SNPs (--min-alleles 2 -- max-alleles 2), minimal distance between the SNPs to reduce linkage=50 kbp (--thin 50000), no missing genotypes and indels (--max-missing-count 0 --remove-indels). Then the VCF files of each family were converted to COLONY format and 1900 random loci were chosen and tested with the following parameters to infer the family relationship: inbreeding absent, diploid species, polygamy for males and females, weak sibship prior, unknown population frequency, one long run, full-likelihood method, very high precision. Based on these results, offspring of each mother were clustered by inferred father, resulting in 1-4 half-sib families per mother. Patterns of variation within and across families were explored with a PCA.

#### 4.2.3 Linkage map construction

The VCF files derived from the genome mapping and half-sib family inference step were filtered allowing minimal phred-scaled variant quality=500 (--minQ 500), minimal genotype quality=10 (--minGQ 10), minor allele count per loci=8 (--mac 8), minimum and maximum alleles per loci=2 (--min-alleles 2 --max-alleles 2), maximum missing count=20 (--max-missing-count 20). We used relatively liberal thresholds for filtering because the rest of the linkage map pipeline was based on genotype-likelihood values instead of the exact genotypes and therefore automatically accounts for error-probabilities.

In the subsequent steps we used the Lep-MAP3 v0.2 pipeline (Rastas 2017) to convert VCF files and produce the linkage map. Initially the filtered VCF file was converted to the Lep-MAP3 posterior probabilities format with the developer’s script *vcf2posterior.awk*. Then we added pedigree information from a custom Excel spreadsheet at the header of the Lep-MAP3 posterior probabilities file and posterior values of zero for the father genotypes, which were absent in our dataset. Next, we imputed the missing father genotypes with the command *ParentCall2* using flags halfSibs=1, familyLimit=0.5, removeNonInformative=1 and outputParentPosterior= 1.

In order to construct the linkage map we used different combinations of parameters to empirically optimise the number of linkage groups and markers in them based on the highest LOD score. Following the Lep-MAP3 pipeline, first we separated linkage groups with the module *SeparateChromosomes2* using LOD score=10.7 (lodLimit=10.7) and at least 150 markers per linkage group. Then we added additional markers to the generated linkage groups using module *JoinSingles2All* with the minimal LOD score=9 (lodLimit=9), minimal difference to assign a marker to one or another linkage group=5 (lodDifference=5), and sequential iterations to assign markers (iterate=1). Finally, the markers were ordered using *OrderMarkers2* module with Kosambi distances (useKosambi=1), separately for each sex (sexAveraged=0) and in the course of eight subsequent iterations (numMergeIterations=8). At the final stage of linkage map construction we arranged the scaffolds into linkage groups using ALLMAPS software (Tang et al. 2015) using both the male and female linkage maps simultaneously.

### 4.3 Assembly QC and validation

In order to validate the assembly and avoid retaining erroneously joined contigs, we used two steps that are based on independent evidence. First, we used REAPR v1.0.18 software (Hunt et al. 2013), which utilizes mate-pair reads mapped to the assembly, to find extensive drops in the fragment coverage along the genome; such signals can be attributed to erroneous contig joinings. To do so we mapped the 8-12 kbp mate-pair library reads to the genome using the SMALT mapper (Ponstingl and Ning 2010) as recommended by the REAPR developers and then identified suspicious regions in the genome. Additionally, we mapped PacBio and RNA-seq reads to the assembly using BWA mem and estimated fragment coverage at those suspicious regions using BEDTools v2.26.0 software (Quinlan 2014). If the PacBio or RNA-seq fragment coverage was less than 2x then we broke the scaffolds in the identified suspicious regions.

Second, we used the *Zootoca vivipara* high-density linkage map to break potential contig misjoins using ALLMAPS v0.7.7 software (Tang et al. 2015). After linkage group construction we applied the ALLMAPS command *split* with at least three significant matches to different linkage groups (--chunk=3), then the coordinates of the potential splits were refined based on the location of gaps in the assembly *(gaps* and *refine* commands), and scaffolds were broken in those coordinates. Then the 500 bp long regions surrounding the RAD-seq markers were extracted from the scaffold assembly (bedtools *getfasta)* and remapped with *blastn* (-max_target_seqs 1 -outfmt 6 -evalue 1e-100) to the broken assembly in order to build the final linkage groups with the ALLMAPS *path* command.

During the course of the genome assembly we explored different assembly methods and pipelines and controlled quality of the output using BUSCO v2.0.1 (Simão et al. 2015; tetrapoda_odb9 database with 3950 single-copy orthologs) and QUAST v4.4 software (Gurevich et al. 2013) quality metrics. BUSCO searches for a set of reliable single-copy orthologues that occur with the highest probability in the given taxonomic group. QUAST estimates different metrics of the genome contiguity and completeness. Metrics were compared at each stage.

### 4.4 Genome annotation

#### 4.4.1 Gene annotation

To annotate the Z. *vivipara* genome we employed homology-based, *ab initio* prediction and RNA-seq based methods, which were combined at the final stage. For the homology-based gene prediction we used GeMoMa 1.4.2 software (Keilwagen et al. 2016), which exploits dynamic programming and intron position conservation supported by RNA-seq data to precisely predict genes in evolutionary related genomes. We aligned the cleaned RNA-seq reads to the *Z. vivipara* scaffolds using STAR v020201 software (Dobin et al. 2103) with the following settings: -- twopass Mode Basic --seedSearchStartLmax 25 and extracted intron positions using GeMoMa *CLI ERE* command. Then we downloaded genomes and annotations of Japanese gecko *Gekko japonicus* (Liu et al. 2015), chicken *Gallus gallus* (International Chicken Genome Sequencing Consortium 2004), and green anole lizard *Anolis carolinensis* (Alföldi et al. 2011) from the NCBI RefSeq (Pruitt et al. 2006) and extracted exons for each species finally converting them into amino acid sequences using GeMoMa *CLI Extractor* command. The translated exons were aligned to *Z. vivipara* scaffolds using *tblastn* (Gertz et al. 2006) in parallel with assistance of the GNU PARALLEL v20161022 (Tange 2011) software. Then, the gene models were identified using *CLI GeMoMa* module with the maximum allowed intron length equal to 50 kbp, the maximum number of transcripts per gene equal to 1 and at least 2 split reads needed to infer an intron border position with RNA-seq data. The gene predictions for each species were further combined and filtered using GeMoMa Annotation Filter *(CLI GAF*) with the following settings: relative score filter =2, common border filter=1 and other settings were set as default.

To invoke RNA-seq evidence for the gene prediction we used StringTie v1.3.1c software (Pertea et al. 2015) to assemble transcripts based on the STAR-aligned BAM files and then filtered and improved them using TACO v0.7.3 (Niknafs et al. 2017) software with the following settings: -- filter-min-expr 0.1, --max-isoforms 1. After, the transcripts were extracted from the scaffolds using *gtf_genome_to_cdna_fasta.pl* script of the TransDecoder v5.0.2 pipeline (https://github.com/TransDecoder/TransDecoder) and coding sequences were identified in them with the TransDecoder.LongOrfs module and then blasted against the Swiss-Prot (Boutet et al. 2016) database using the DIAMOND v0.9.13 software (Buchfink et al. 2015) with the following settings: -c 1 -k 1 -p 60 -e 1e-5 --more-sensitive. Finally, we retained at most one open reading frame (ORF) per transcript using the TransDecoder.Predict module and hints from the DIAMOND *blastp* output and combined them with the the TACO output to extract protein-coding gene models with RNA-seq evidence and homology to existing proteins with the script *cdna_alignment_orf_to_genome_orf.pl*.

The *de novo* annotation part was accomplished with the AUGUSTUS v3.3 software (Stanke et al. 2003) using scaffolds with repeats masked by WindowMasker v2.7.0 (Morgulis et al. 2005) and empirically optimised Hidden Markov Model from the BUSCO output (--species).

At the final stage the three annotations were combined with the EVidenceModeler v1.1.1 software (Haas et al. 2008) during two stages. First, the consensus gene-models were calculated and extracted. Then, the consensus proteins were blasted against the Swiss-Prot (Boutet et al. 2016) database using the DIAMOND v0.9.13 software (Buchfink et al. 2015) with the following settings: -c 1 -k 1 -p 28 -e 1e-5 --more-sensitive and genes without any homology to the database were filtered out.

## 5 ACKNOWLEDGEMENTS

We are grateful for the help in library preparation and RNA extraction to Aileen Adams and Elizabeth Kilbride. We thank NBAF Edinburgh (especially Karim Gharbi and Helen Gunter) and NBAF Liverpool for advice, preparations, and sequencing. This research was funded by Natural Environment Research Council (NE/N003942/1, NBAF964, and NBAF1018).

## 6 USE OF ANIMALS

The genome lizard is a female (heterogametic sex) from Isle of Cumbrae, Scotland, collected for this study in 2015 with permission of Scottish Natural Heritage (64972). Her name is Vivacia. Samples for linkage mapping were collected and sampled non-lethally before release, with permission from Bezirkshauptmannschaft Hermagor, Austria (HE3-NS-959/2013).

## 7 AVAILABILITY OF DATA AND MATERIALS

The genome assembly and annotation are freely available from authors by request.

## REFERENCES

Alföldi, J., Di Palma, F., Grabherr, M., Williams, C., Kong, L., Mauceli, E., … & Ray, D. A. (2011). The genome of the green anole lizard and a comparative analysis with birds and mammals. Nature, 477, 587.

Andrade, P., Pinho, C., i de Lanuza, G. P., Afonso, S., Brejcha, J., Rubin, C. J., … & Pellitteri-Rosa, D. (2018). Regulatory changes in pterin and carotenoid genes underlie balanced color polymorphisms in the wall lizard. bioRxiv, 481895.

Andrews, S. (2010). FastQC. A quality control tool for high throughput sequence data.

Blackburn, D. G. (2006). Squamate reptiles as model organisms for the evolution of viviparity. Herpetological Monographs, 20, 131–146.

Bolger, A. M., Lohse, M., & Usadel, B. (2014). Trimmomatic: a flexible trimmer for Illumina sequence data. Bioinformatics, 30, 2114–2120.

Boutet, E., Lieberherr, D., Tognolli, M., Schneider, M., Bansal, P., Bridge, A. J., … & Xenarios, I. (2016). UniProtKB/Swiss-Prot, the manually annotated section of the UniProt KnowledgeBase: how to use the entry view. In Plant Bioinformatics (pp. 23–54). Humana Press, New York, NY.

Buchfink, B., Xie, C., & Huson, D. H. (2015). Fast and sensitive protein alignment using DIAMOND. Nature Methods, 12, 59.

Castoe, T. A., De Koning, A. J., Hall, K. T., Card, D. C., Schield, D. R., Fujita, M. K., … & Reyes-Velasco, J. (2013). The Burmese python genome reveals the molecular basis for extreme adaptation in snakes. Proceedings of the National Academy of Sciences, 110, 20645–20650.

Catchen, J. M., Amores, A., Hohenlohe, P., Cresko, W., & Postlethwait, J. H. (2011). Stacks: building and genotyping loci de novo from short-read sequences. G3: Genes, Genomes, Genetics, 1, 171–182.

Deakin, J. E., & Ezaz, T. (2014). Tracing the evolution of amniote chromosomes. Chromosoma, 123, 201–216.

Desmet, W. (1981). The nuclear Feulgen-DNA content of the vertebrates (especially reptiles), as measured by fluorescence cytophotometry, with notes on the cell and the chromosome size. Acta Zoologica et Pathologica Antverpiensia, 76, 119–167.

Dobin, A., Davis, C. A., Schlesinger, F., Drenkow, J., Zaleski, C., Jha, S., … & Gingeras, T. R. (2013). STAR: ultrafast universal RNA-seq aligner. Bioinformatics, 29, 15–21.

English, A. C., Richards, S., Han, Y., Wang, M., Vee, V., Qu, J., … & Gibbs, R. A. (2012). Mind the gap: upgrading genomes with Pacific Biosciences RS long-read sequencing technology. PloS one, 7, e47768.

Gao, S., Bertrand, D., Chia, B. K., & Nagarajan, N. (2016). OPERA-LG: efficient and exact scaffolding of large, repeat-rich eukaryotic genomes with performance guarantees. Genome Biology, 17, 102.

Gertz, E. M., Yu, Y. K., Agarwala, R., Schäffer, A. A., & Altschul, S. F. (2006). Composition-based statistics and translated nucleotide searches: improving the TBLASTN module of BLAST. BMC Biology, 4, 41.

Guarnizo, C. E., Werneck, F. P., Giugliano, L. G., Santos, M. G., Fenker, J., Sousa, L., … & Dorado-Rodrigues, T. F. (2016). Cryptic lineages and diversification of an endemic anole lizard (Squamata, Dactyloidae) of the Cerrado hotspot. Molecular Phylogenetics and Evolution, 94, 279–289.

Gunter, H. M., Fan, S., Xiong, F., Franchini, P., Fruciano, C., & Meyer, A. (2013). Shaping development through mechanical strain: the transcriptional basis of diet-induced phenotypic plasticity in a cichlid fish. Molecular Ecology, 22, 4516–4531.

Gurevich, A., Saveliev, V., Vyahhi, N., & Tesler, G. (2013). QUAST: quality assessment tool for genome assemblies. Bioinformatics, 29, 1072–1075.

Haas, B. J., Salzberg, S. L., Zhu, W., Pertea, M., Allen, J. E., Orvis, J., … & Wortman, J. R. (2008). Automated eukaryotic gene structure annotation using EVidenceModeler and the Program to Assemble Spliced Alignments. Genome Biology, 9, 1.

Horreo, J. L., Pelaez, M. L., Suarez, T., Breedveld, M. C., Heulin, B., Surget-Groba, Y., . . . Fitze, P. (2018). Phylogeography, evolutionary history and effects of glaciations in a species (Zootoca vivipara) inhabiting multiple biogeographic regions. Journal of Biogeography, 45, 1616–1627.

Hunt, M., Kikuchi, T., Sanders, M., Newbold, C., Berriman, M., & Otto, T. D. (2013). REAPR: a universal tool for genome assembly evaluation. Genome Biology, 14, R47.

International Chicken Genome Sequencing Consortium. (2004). Sequence and comparative analysis of the chicken genome provide unique perspectives on vertebrate evolution. Nature, 432, 695.

Irisarri, I., Baurain, D., Brinkmann, H., Delsuc, F., Sire, J. Y., Kupfer, A., … & Philippe, H. (2017). Phylotranscriptomic consolidation of the jawed vertebrate timetree. Nature Ecology and Evolution, 1, 1370.

Jones, O. R., & Wang, J. (2010). COLONY: a program for parentage and sibship inference from multilocus genotype data. Molecular Ecology Resources, 10, 551–555.

Kajitani, R., Toshimoto, K., Noguchi, H., Toyoda, A., Ogura, Y., Okuno, M., … & Kohara, Y. (2014). Efficient de novo assembly of highly heterozygous genomes from whole-genome shotgun short reads. Genome Research, 24, 1384–1395.

Keilwagen, J., Wenk, M., Erickson, J. L., Schattat, M. H., Grau, J., & Hartung, F. (2016). Using intron position conservation for homology-based gene prediction. Nucleic Acids Research, 44, e89–e89.

King, B., & Lee, M. S. (2015). Ancestral state reconstruction, rate heterogeneity, and the evolution of reptile viviparity. Systematic Biology, 64, 532–544.

Kolora, S. R. R., Weigert, A., Saffari, A., Kehr, S., Costa, M. B. W., Spröer, C., … & Overmann, J. (2018). Divergent evolution in the genomes of closely-related lacertids, Lacerta viridis and L. bilineata and implications for speciation. GigaScience, giy160.

Koren, S., Walenz, B. P., Berlin, K., Miller, J. R., Bergman, N. H., & Phillippy, A. M. (2017). Canu: scalable and accurate long-read assembly via adaptive k-mer weighting and repeat separation. Genome Research, 27, 722–736.

Kupriyanova, L., Kuksin, A., & Odierna, G. (2008). Karyotype, chromosome structure, reproductive modalities of three Southern Eurasian populations of the common lacertid lizard, *Zootoca vivipara* (Jacquin, 1787). Acta Herpetologica, 3, 99–106.

Kupriyanova, L., Niskanen, M., & Oksanen, T. A. (2014). Karyotype dispersal of the common lizard Zootoca vivipara (Lichtenstein, 1823) in eastern and northeastern Fennoscandia. Memoranda Societatis pro Fauna et Flora Fennica.

Laloi, D., M. Richard, J. Lecomte, M. Massot, and J. Clobert. 2004. Multiple paternity in clutches of common lizard Lacerta vivipara: data from microsatellite markers. Molecular Ecology 13, 719–23.

Li, H., & Durbin, R. (2009). Fast and accurate short read alignment with Burrows-Wheeler transform. Bioinformatics, 25, 1754–1760.

Liu, Y., Zhou, Q., Wang, Y., Luo, L., Yang, J., Yang, L., … & Li, M. (2015). Gekko japonicus genome reveals evolution of adhesive toe pads and tail regeneration. Nature Communications, 6, 10033.

Luo, R., Liu, B., Xie, Y., Li, Z., Huang, W., Yuan, J., … & Tang, J. (2012). SOAPdenovo2: an empirically improved memory-efficient short-read de novo assembler. Gigascience, 1, 18.

Marçais, G., Yorke, J. A., & Zimin, A. (2015). QuorUM: an error corrector for Illumina reads. PLoS One, 10, e0130821.

Morgulis, A., Gertz, E. M., Schäffer, A. A., & Agarwala, R. (2005). WindowMasker: window-based masker for sequenced genomes. Bioinformatics, 22, 134–141.

Neaves, W. B., & Baumann, P. (2011). Unisexual reproduction among vertebrates. Trends in Genetics, 27, 81–88.

Niknafs, Y. S., Pandian, B., Iyer, H. K., Chinnaiyan, A. M., & Iyer, M. K. (2017). TACO produces robust multisample transcriptome assemblies from RNA-seq. Nature methods, 14, 68.

O’Connell, J., Schulz-Trieglaff, O., Carlson, E., Hims, M. M., Gormley, N. A., & Cox, A. J. (2015). NxTrim: optimized trimming of Illumina mate pair reads. Bioinformatics, 31, 2035–2037.

Odierna, G., Aprea, G., Capriglione, T., & Puky, M. (2004). Chromosomal evidence for the double origin of viviparity in the European common lizard, Lacerta (Zootoca) vivipara. Herpetological Journal, 14, 157–160.

Oliver, P. M., Adams, M., Lee, M. S., Hutchinson, M. N., & Doughty, P. (2009). Cryptic diversity in vertebrates: molecular data double estimates of species diversity in a radiation of Australian lizards (Diplodactylus, Gekkota). Proceedings of the Royal Society of London B: Biological Sciences, 276, 2001–2007.

Pertea, M., Pertea, G. M., Antonescu, C. M., Chang, T. C., Mendell, J. T., & Salzberg, S. L. (2015). StringTie enables improved reconstruction of a transcriptome from RNA-seq reads. Nature Biotechnology, 33, 290.

Peterson, B. K., Weber, J. N., Kay, E. H., Fisher, H. S., & Hoekstra, H. E. (2012). Double digest RADseq: an inexpensive method for de novo SNP discovery and genotyping in model and non-model species. PloS one, 7, e37135.

Pinho, C., Harris, D. J., & Ferrand, N. (2007). Comparing patterns of nuclear and mitochondrial divergence in a cryptic species complex: the case of Iberian and North African wall lizards (Podarcis, Lacertidae). Biological Journal of the Linnean Society, 91, 121–133.

Ponstingl, H., & Ning, Z. (2010). SMALT-a new mapper for DNA sequencing reads. F1000 Posters, 1, 313.

Pruitt, K. D., Tatusova, T., & Maglott, D. R. (2006). NCBI reference sequences (RefSeq): a curated non-redundant sequence database of genomes, transcripts and proteins. Nucleic Acids Research, 35, D61–D65.

Pyron, R. A., & Burbrink, F. T. (2014). Early origin of viviparity and multiple reversions to oviparity in squamate reptiles. Ecology Letters, 17, 13–21.

Quinlan, A. R. (2014). BEDTools: the Swiss-army tool for genome feature analysis. Current Protocols in Bioinformatics, 47, 11.12.1–11.12.34.

Rastas, P. (2017). Lep-MAP3: robust linkage mapping even for low-coverage whole genome sequencing data. Bioinformatics, 33, 3726–3732.

Recknagel, H., Jacobs, A., Herzyk, P., & Elmer, K. R. (2015). Double-digest RAD sequencing using Ion Proton semiconductor platform (ddRADseq-ion) with nonmodel organisms. Molecular Ecology Resources, 15, 1316–1329.

Recknagel, H., Kamenos, N., & Elmer, K. R. (2018). Common lizards break Dollo’s law of irreversibility: Genome-wide phylogenomics support a single origin of viviparity and re-evolution of oviparity. Molecular Phylogenetics and Evolution, 127, 579–588

Sambrook, J., & Russell, D. W. (2006). Purification of nucleic acids by extraction with phenol: chloroform. Cold Spring Harbor Protocols, 2006, pdb–prot4455.

Simão, F. A., Waterhouse, R. M., Ioannidis, P., Kriventseva, E. V., & Zdobnov, E. M. (2015). BUSCO: assessing genome assembly and annotation completeness with single-copy orthologs. Bioinformatics, 31, 3210–3212.

Simpson, J. T. (2014). Exploring genome characteristics and sequence quality without a reference. Bioinformatics, 30, 1228–1235.

Sites Jr, J. W., Reeder, T. W., & Wiens, J. J. (2011). Phylogenetic insights on evolutionary novelties in lizards and snakes: sex, birth, bodies, niches, and venom. Annual Review of Ecology, Evolution, and Systematics, 42, 227–244.

Stanke, M., & Waack, S. (2003). Gene prediction with a hidden Markov model and a new intron submodel. Bioinformatics, 19, ii215–ii225.

Stanke, M., Steinkamp, R., Waack, S., & Morgenstern, B. (2004). AUGUSTUS: a web server for gene finding in eukaryotes. Nucleic Acids Research, 32, W309–W312.

Surget-Groba, Y., Heulin, B., Guillaume, C. P., Thorpe, R. S., Kupriyanova, L., Vogrin, N., … & Odierna, G. (2001). Intraspecific phylogeography of Lacerta vivipara and the evolution of viviparity. Molecular Phylogenetics and Evolution, 18, 449–459.

Tang, H., Zhang, X., Miao, C., Zhang, J., Ming, R., Schnable, J. C., … & Lu, J. (2015). ALLMAPS: robust scaffold ordering based on multiple maps. Genome Biology, 16, 3.

Tange, O. (2011). Gnu parallel-the command-line power tool. The USENIX Magazine, 36, 42–47.

Tollis, M., Hutchins, E. D., Stapley, J., Rupp, S. M., Eckalbar, W. L., Maayan, I., … & May, C. M. (2018). Comparative Genomics Reveals Accelerated Evolution in Conserved Pathways during the Diversification of Anole Lizards. Genome Biology and Evolution, 10, 489–506.

Uetz, P., Hošek, J. (Eds.), 2015. The Reptile Database. <http://www.reptile-database.org> (accessed 18.02.15).

Uller, T., & Olsson, M. (2008). Multiple paternity in reptiles: patterns and processes. Molecular Ecology, 17, 2566–2580.

Vercken, E., Massot, M., Sinervo, B., & Clobert, J. (2007). Colour variation and alternative reproductive strategies in females of the common lizard Lacerta vivipara. Journal of Evolutionary Biology, 20, 221–232.

Vinogradov, A. E. (1998). Genome size and GC-percent in vertebrates as determined by flow cytometry: The triangular relationship. Cytometry Part A, 31, 100–109.

Zhang, S. V., Zhuo, L., & Hahn, M. W. (2016). AGOUTI: improving genome assembly and annotation using transcriptome data. GigaScience, 5, 31.

Zheng, Y., & Wiens, J. J. (2016). Combining phylogenomic and supermatrix approaches, and a time-calibrated phylogeny for squamate reptiles (lizards and snakes) based on 52 genes and 4162 species. Molecular Phylogenetics and Evolution, 94, 537–547.

